# Helical Fibrillar Microstructure of Tendon using Serial Block-Face SEM and a Mechanical Model for Interfibrillar Load Transfer

**DOI:** 10.1101/547281

**Authors:** Babak N. Safa, John M. Peloquin, Jessica R. Natriello, Jeffrey L. Caplan, Dawn M. Elliott

## Abstract

Tendon’s hierarchical structure allows for load transfer between its fibrillar elements at multiple length scales. Tendon microstructure is particularly important, because it includes the cells and their surrounding collagen fibrils, where mechanical interactions can have potentially important physiological and pathological contributions. However, the three-dimensional microstructure and the mechanisms of load transfer in that length scale are not known. It has been postulated that interfibrillar matrix shear or direct load transfer via the fusion/branching of small fibrils are responsible for load transfer, but the significance of these mechanisms is still unclear. Alternatively, the helical fibrils that occur at the microstructural scale in tendon may also mediate load transfer, however, these structures are not well studied due to the lack of a three-dimensional visualization of tendon microstructure. In this study, we used serial block-face scanning electron microscopy (SBF-SEM) to investigate the three-dimensional microstructure of fibrils in rat tail tendon. We found that tendon fibrils have a complex architecture with many helically wrapped fibrils. We studied the mechanical implications of these helical structures using finite element modeling and found that frictional contact between helical fibrils can induce load transfer even in the absence of matrix bonding or fibril fusion/branching. This study is significant in that it provides a three-dimensional view of the tendon microstructure and suggests friction between helically wrapped fibrils as a mechanism for load transfer, which is an important aspect of tendon biomechanics.

## Introduction

Tendon’s hierarchical structure allows for load transfer between its fibrillar elements across multiple length scales (Kastelic et al., 1978; Pensalfini et al., 2014; Screen et al., 2005), which results in remarkable capabilities to withstand stress and endure repetitive loading (Snedeker and Foolen, 2017). The microscale structure and function are particularly important, because this is the scale where the cells and their surrounding collagen fibrils interface, and these mechanical interactions can have important physiological and pathological contributions. In particular, there is evidence of microscale sliding and shear load transfer that is highly likely to represent sliding between fibrils at small strains (less than 2%) and at larger strains this microscale sliding is nonrecoverable, indicating tissue damage (Lee et al., 2017; Screen et al., 2005; Szczesny and Elliott, 2014a). However, the underlying mechanisms of load transfer between tendon fibrils are still unknown. Tendon fibrils are collagenous structures (diameter ∼100 nm) that are the building blocks of tendon microstructure (Kannus, 2000). The fibrils are responsible for supporting external mechanical loading. Interfibrillar matrix molecules such as glycosaminoglycan chains (GAG) have been postulated to be responsible for load transfer between fibrils (Ahmadzadeh et al., 2013; Redaelli et al., 2003); however, removal of a wide range of extracellular matrix components, including GAG, does not affect the mechanical response (Szczesny et al., 2017), or has minimal consequences (Fessel and Snedeker, 2009). It is not likely that the load transfer in tendon is solely, or majorly, mediated via interfibrillar matrix. Thus, the microscale architecture of the collagen fibrous network itself is likely to have a role in mediating load transfer.

Serial block-face scanning electron microscopy (SBF-SEM) makes it possible to visualize the 3D microscale architecture of tissue in great detail, and to reveal the fibrillar architecture with several mechanical implications (Pingel et al., 2014; Starborg et al., 2013). SBF-SEM is an advanced electron microscopy technique that takes sequential SEM images of the cross-section of tissue; these images combine to provide a three-dimensional view of the microstructure (Hashimoto et al., 2016; Starborg et al., 2013). Using SBF-SEM with a short scan depth (8.7 µm), our group showed that there is a small fibril angular dispersion and that the fusion/branching of small fibrils might be responsible for interfibrillar load transfer (Szczesny et al., 2017, 2014). Longer scan depths (∼100 µm) showed that fibril fusion/branching and fibril ends also exist in tendons (Svensson et al., 2017), and that helical fibril patterns form in microscale during tendon development in juvenile tail tendons (Kalson et al., 2015; Starborg and Kadler, 2015). The existence of helical fibrils in collagen microscale and nanoscale was shown using two-dimensional light, atomic force, electron, and X-ray scattering microscopy in tendons and other collagenous tissues (Altraud et al., 1987; Bozec et al., 2007; Folkhard et al., 1987; Franchi et al., 2010; Kalson et al., 2012; Lillie et al., 1977; Orgel et al., 2006; Reed et al., 1956; Vidal, 2003); however, these two-dimensional visualizations do not provide the three-dimensional structure.

Experimental observations and finite element (FE) modeling suggest that helical structures in tendon may have significant mechanical effects. The rotation and high Poisson’s ratio observed during tendon’s axial loading have been attributed to such structures (Buchanan et al., 2017; Reese et al., 2010; Thorpe et al., 2013). Furthermore, some FE models have studied the groups of helical fibrils by combining the fibrils with interfibrillar matrix in a mesh to produce the nonlinear stress response of fascicles (Carniel et al., 2019; Reese et al., 2010). Despite the potential of helical fibril organization to affect tendon’s microscale mechanics, little is known about individual groups of helically wrapped fibrils and their mechanical implications. We hypothesized that helical wrapping can induce frictional load transfer between fibrils, allowing for mechanical interfibrillar load transfer without an intermediate matrix. This would imply that friction between helically wrapped fibrils can contribute to load transfer, in addition to interfibrillar matrix shear and fibril fusion/branching.

The scope of this contribution was to visualize the three-dimensional fibril organization of rat tail tendon and to study the potential of interfibrillar friction within helically wrapped groups of fibrils to serve as a mechanism for load transfer. First, we visualized the microstructure of tendon in three dimensions using SBF-SEM. We found a complex network, with many helically wrapped fibrils. These observations informed the second part of this study, in which we used finite element (FE) analysis to test the hypothesis that frictional contact between helically wrapped fibrils can transfer stress (load) between fibrils without a need for a mediating matrix. This study elucidates new aspects of tendon microstructure, providing a detailed image of fibril tortuosity, fusion/branching, and organization into helical groups. Importantly, our results establish interfibrillar friction as a new mechanism for interfibrillar load transfer, among the several proposed load transfer mechanisms in the field, advancing our knowledge about microscale structure–mechanics relationships.

## Methods

### Serial block-face SEM imaging

A tail tendon fascicle from a three-month-old male Sprague-Dawley rat was dissected as previously described (Safa et al., 2017), and used for SBF-SEM imaging. To prepare for imaging, the fascicle was placed in PBS for 8 hours to equilibrate and then soaked overnight at 4°C in a solution of 2% glutaraldehyde and 2% paraformaldehyde in 0.1 M sodium cacodylate buffer. The sample was subsequently stained and resin-embedded, according to established techniques (Starborg et al., 2013).

The transverse cross-section was scanned in series under 1.78 kV in low-vacuum pressure with three microseconds dwelling time using an Apreo VolumeScope (Thermo Fisher Scientific, Waltham MA). After the end of the scanning of each section, a 200 nm layer of the block’s face was removed by means of a mechanical slicer and the scanning was repeated. The in-plane resolution of the scans was 10 nm/pixel with 432 slices that covers a total volume of (20.27 µm × 16.74 µm × 86.20 µm), where the third dimension is measured along the fascicle’s axial direction.

### Segmentation and data analysis

The SEM images were smoothed using a Gaussian blur filter and a representative subset of fibrils were manually segmented through the entire 3D image stack (Fig. 1A). Only fibrils that spanned the entire image stack, as many fibrils did not entirely fit in the scanned volume or some fibrils were otherwise discontinuous, were used for analysis (*n* = 42 fibrils). The segmentation was done using Seg3D (seg3d.org). Since the axial distance between subsequent images was small (200 nm) the fibrils appear as semi-circular spots that move in-plane allowing for the fibrils to be inspected and tracked through the image stack (Fig. 1A-C).

**Figure 1:**
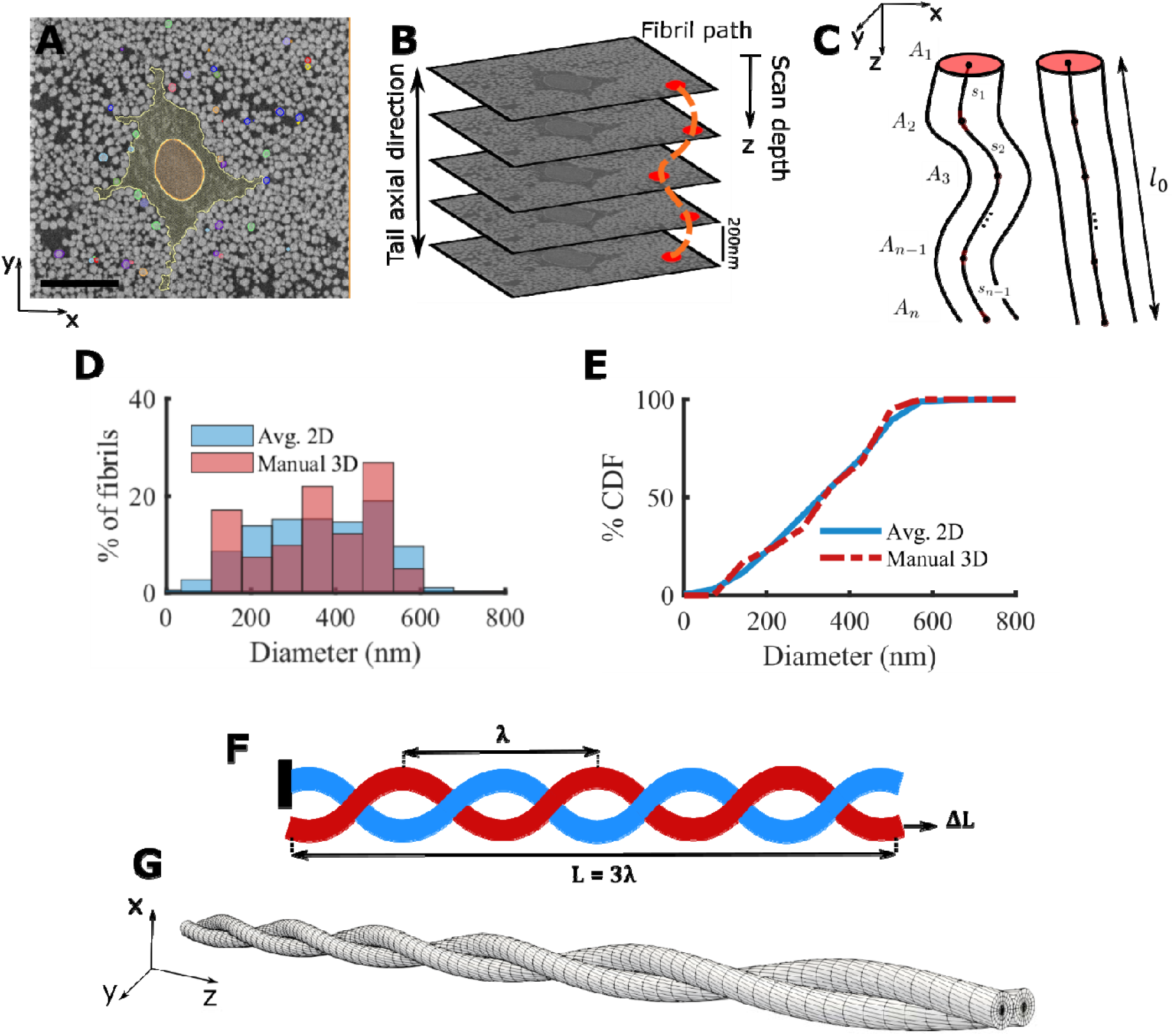
(A) Representative SEM image and in-plane segmentation (scale bar = 5 *µ*m), (B) SBF-SEM imaging process showing sequential stacks of images and a schematic fibril label (dotted orange line) along the tendon length, (C) schematics of fibrils with large (left) and low (right) tortuosity that shows the cross-sectional area (*A*_*i*_) of fibrils throughout the scan depth and the incremental distance (*s*_*i*_) between the center of the fibrils. (D) By comparing the distribution and (E) cumulative distribution function (CDF) of the diameter using the averaged 2D automatic segmentation of all fibrils (blue) and the 3D manual segmentation of a subset of 42 fibrils (red), it is evident that the manually segmented fibrils are a representative of the full fibril population. (F) For the finite element analysis a model with three full turns is used (L = 3A). The boundary condition is that one fibril (blue) is anchored (left), while the other fibril (red) is pulled in the axial direction (arrow, right) to a deformation of Δ*L* = *Lε*. (G) The mesh used for the finite element simulations shows that the fibrils were initially in contact throughout the length.

To confirm that the manually segmented fibrils were a representative selection of all the fibrils, we segmented all fibrils in each of ten equally spaced 2D sections by thresholding (ImageJ, imagej.nih.gov). The distribution of fibril diameter was calculated and compared to that of the manual 3D fibril segmentations (Fig. 1D and E). The distributions matched, confirming that the manually segmented fibrils were a representative sample. The manual 3D fibril segmentations were used for all subsequent analysis.

To determine the variation in fibril diameter along the length, we calculated the normalized diameter of each fibril in each section in the image stack. The normalized diameter for a fibril at section *i* is defined as

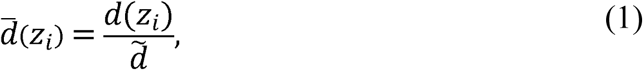

where *z*_*i*_ is the scan depth, 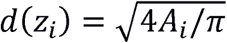 is the diameter of the fibril, *A*_*i*_ is the cross-sectional area of the fibril, and 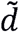 is the median of *d*(*z*_*i*_) across all sections (Fig. 1C). To quantify the complexity of the fibrillar network, we calculated the percent tortuosity for each fibril as

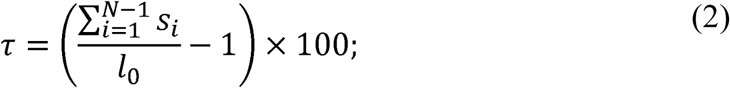

here, *l*_0_ is the end-to-end distance of the fibrils, and *s*_*i*_ is the distance between fibril centroids in adjacent sections, and *N* is the number of sections (Fig. 1C). Approximately half of the segmented fibrils (*n* = 20) were in helical groups; for these fibrils we calculated the pitch (*λ*) by diving the total scan depth (86.20 µm) by the number of turns in the helices.

### Statistics

The variation in diameter along the length was tested by calculating, for each section, the 95% confidence interval of normalized diameter (Eq. 1) across the fibrils in that section. To test the correlation between fibril diameter and tortuosity, we conducted a linear regression analysis between 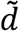 and *τ* for the fibrils in the helical groups and others, separately, using Pearson’s correlation method. Statistical significance was defined as *p* < 0.05.

### Finite element analysis of helical fibril structures

Helical wrapping of fibrils around each other was a commonly observed feature, and we hypothesized that this helical wrapping may provide interfibrillar load transfer. To study this, we developed a three-dimensional finite element (FE) model of a pair of helical fibrils in contact (Fig. 1F and G) using FEBio software (FEBio2.8 febio.org) (Maas et al., 2012). The fibril diameter was taken to be 200 nm, and according to our observations from SBF-SEM, the helix pitch was set at 40 μm. The entire model included three full fibril revolutions (i.e., *L* = 3*λ*, where *L* is the total length). The boundary condition was set as that each fibril had one free-end. In one fibril, the opposite end was anchored; in the other fibril, it was set to move to create 8% axial strain (Fig. 1F). The value of 8% was selected from experimental data as the maximum tissue strain that fibrils may experience prior to failure (Lee et al., 2017).

We used an isotropic compressible neo-Hookean constitutive relation for the fibrils (Bonet and Wood, 1997; Maas et al., 2019):

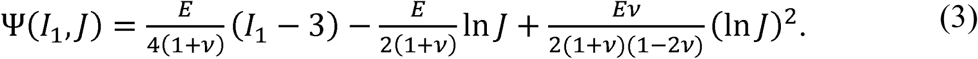

Where, Ψ is the strain energy, *I*_1_ is the first invariant of the right Cauchy-Green strain tensor and *J* is the Jacobian of deformation; *E* is the Young’s modulus, and *v* is the Poisson’s ratio. The model parameters were set as *E* = 1 GPa, *v* = 0.2. The friction parameter between fibrils was set as *µ* = 0.5, and the helix pitch was set as *λ* = 40 µm.

To solve the frictional contact problem between the fibrils we employed the penalty method regularized with an augmented Lagrangian scheme that has been implemented in FEBio (Zimmerman and Ateshian, 2018). In the initial configuration the fibrils were in contact along the entire length. We used a mesh consisting of 20,160 hexahedral trilinear elements and 23,478 nodes based on a mesh sensitivity analysis (Fig 1G). To improve the stability of the contact algorithm we used a frictionless external cylindrical sheath with a weak modulus (10% of fibril modulus) to prevent separation in the intermediate steps of the iterative FE solver. The sheath was highly effective in increasing the stability of the model, and did not alter the final solutions. This was confirmed by using different moduli for the sheath between 0.1% and 1000% of the fibril modulus; the same mechanical response was produced in each case.

The reaction force at the anchored end of the fixed fibril was divided by the cross sectional area and used as the measure for load transfer. To assess the sensitivity of the load transfer, we performed a one-at-a-time parametric sensitivity analysis by varying the model parameters in a range according to the reported values in the literature for fibril properties that are summarize in Table 1 (Chung et al., 2013; Gautieri et al., 2011; Liu et al., 2016; Minary-Jolandan and Yu, 2009; Szczesny and Elliott, 2014a; Wells et al., 2015; Wenger et al., 2007; Yang et al., 2012). Since the experimental data for frictional coefficient of fibril-on-fibril is not available, we used an estimated range (0 < *µ* < 2) based on AFM indentation tests (Chung et al., 2013). We studied the following cases by changing one parameter at a time: *E* = 0.1 to10 GPa,*v* = 0 to 0.4, *µ* = 0 to 2, *λ* = 20 to 80 µm.

**Table 1:**
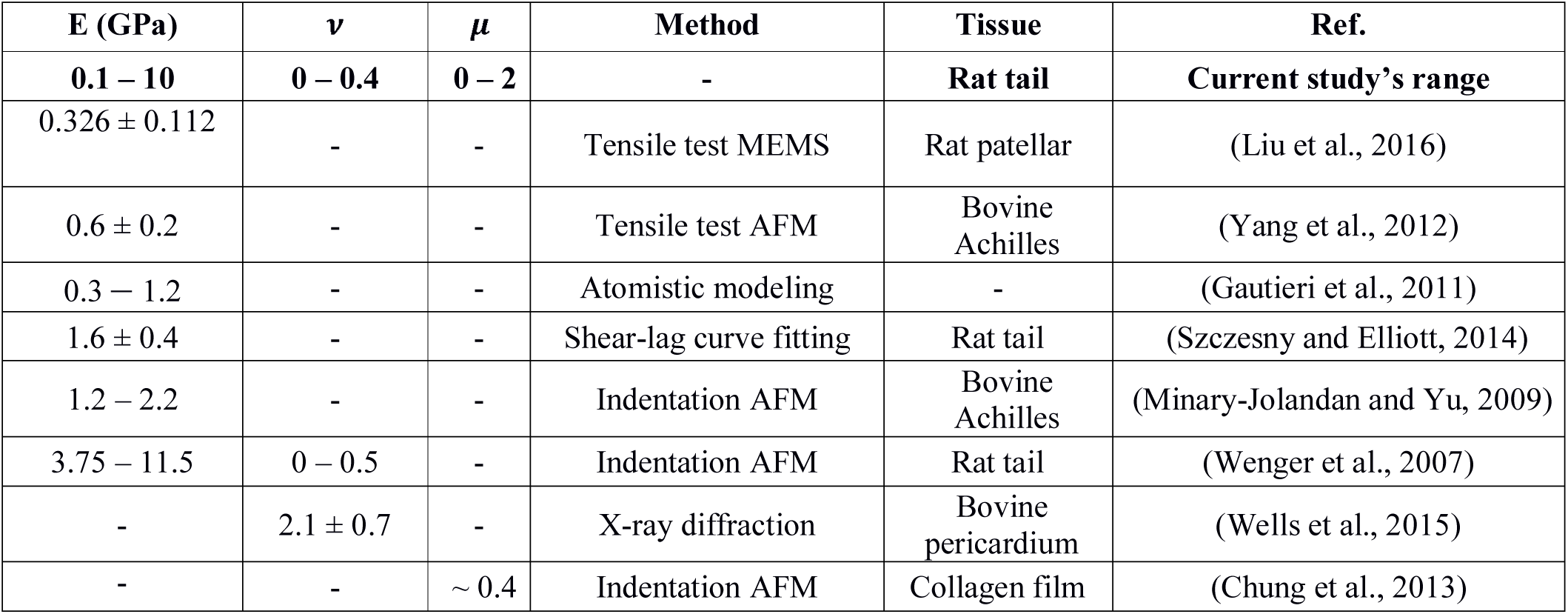
Collagen fibril mechanical properties: Young’s modulus (*E*), Poisson’s ratio (*v*), frictional coefficient (*µ*) in tendon unless otherwise noted.

| **E (GPa)** | **$\nu$** | **$\mu$** | **Method** | **Tissue** | **Ref.** |
| --- | --- | --- | --- | --- | --- |
| **0.1 – 10** | **0 – 0.4** | **0 – 2** | **-** | **Rat tail** | **Current study's range** |
| $0.326 \pm 0.112$ | - | - | Tensile test MEMS | Rat patellar | (Liu et al., 2016) |
| $0.6 \pm 0.2$ | - | - | Tensile test AFM | Bovine Achilles | (Yang et al., 2012) |
| 0.3 – 1.2 | - | - | Atomistic modeling | - | (Gautieri et al., 2011) |
| $1.6 \pm 0.4$ | - | - | Shear-lag curve fitting | Rat tail | (Szczesny and Elliott, 2014) |
| 1.2 – 2.2 | - | - | Indentation AFM | Bovine Achilles | (Minary-Jolandan and Yu, 2009) |
| 3.75 – 11.5 | 0 – 0.5 | - | Indentation AFM | Rat tail | (Wenger et al., 2007) |
| - | $2.1 \pm 0.7$ | - | X-ray diffraction | Bovine pericardium | (Wells et al., 2015) |
| - | - | $\sim 0.4$ | Indentation AFM | Collagen film | (Chung et al., 2013) |

To evaluate the spatial distribution of the induced stress and deformation in the fibrils we plotted axial stress and we also plotted the normalized displacement (*ū*) relative to the mid-section of each fibril along the axial direction. We defined normalized displacement as

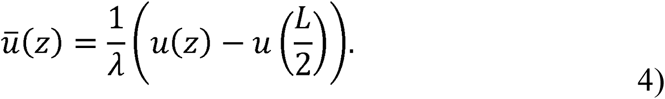

Here, *u*(*z*) is the axial displacement at position *z* along the fibril length.

## Results

### Microstructure of fibrils and SBF-SEM

To describe the microstructure of tendon, we segmented fibrils from the SBF-SEM images in three-dimensions. The segmentation indicated that although the fibrils are mostly axially-aligned, they create a complex network around the cells (Fig. 2A). From the manually segmented fibrils we quantified the fibril diameter throughout the scan depth (Eq. 1) and the percent tortuosity (Eq. 2). As expected, the fibrils’ diameter did not vary along the fibril length (Fig. 2B), which is consistent with previous findings (Svensson et al., 2017). Note that the normalized fibril diameter (Eq. 1) was not significantly different than one across the scan depth (Fig 2B). This observation supports the use of a single median diameter value assigned to each fibril for correlation with the tortuosity. We quantified the % tortuosity (*τ*) of each segmented fibril (Eq. 2), and calculated its correlation with the median diameter of the fibrils. This showed that for all of the fibrils, *τ* was small (< 1%) and for the fibrils in helical groups tortuosity was correlated with diameter (*r* = −0.59, *p* < 0.05), where for the other fibrils it was the same across the diameter sizes (*r* = 0.12, *p* = 0.616) (Fig. 2C). As a result, the smaller fibrils in helical groups are more likely to have higher tortuosity, and thus have a more complex structure. For the tortuosity analysis, one fibril that made a right-angle turn and passed through the cell membrane (Fig. 3C, discussed below) was not included in the correlation.

**Figure 2:**
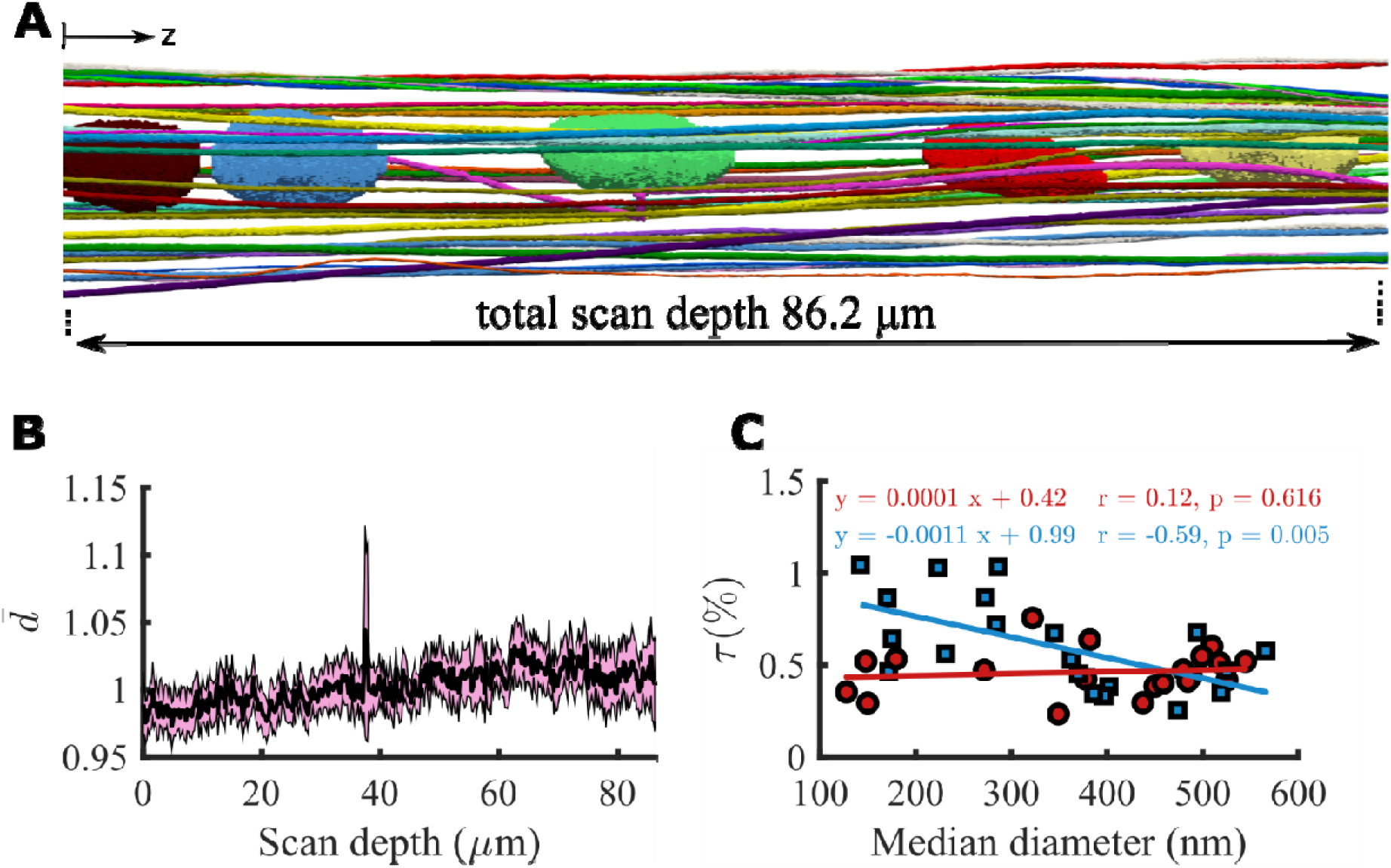
(A) Lateral view of the three dimensional segmentation along the scan length with cell nuclei. Although the fibrils are primarily axially oriented, they have a complex three-dimensional network. (B) the 95% confidence interval (highlighted in pink) of the normalized diameter (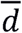, Eq. 1) contains the unity value throughout the scan depth with less than 5% variation, which indicates that the diameter of the fibrils does not change along the scan depth. (C) Percent-tortuosity (*τ*, Eq. 2) was calculated as a measure of the complexity of the fibrillar structure for the fibrils in helical groups (marked with blue squares) and the other fibrils (marked with red circles) separately. The helical fibril tortuosity was correlated with the diameter, with the small diameter fibrils being more tortuous, however, tortuosity was uniform for the other fibrils.

**Figure 3:**
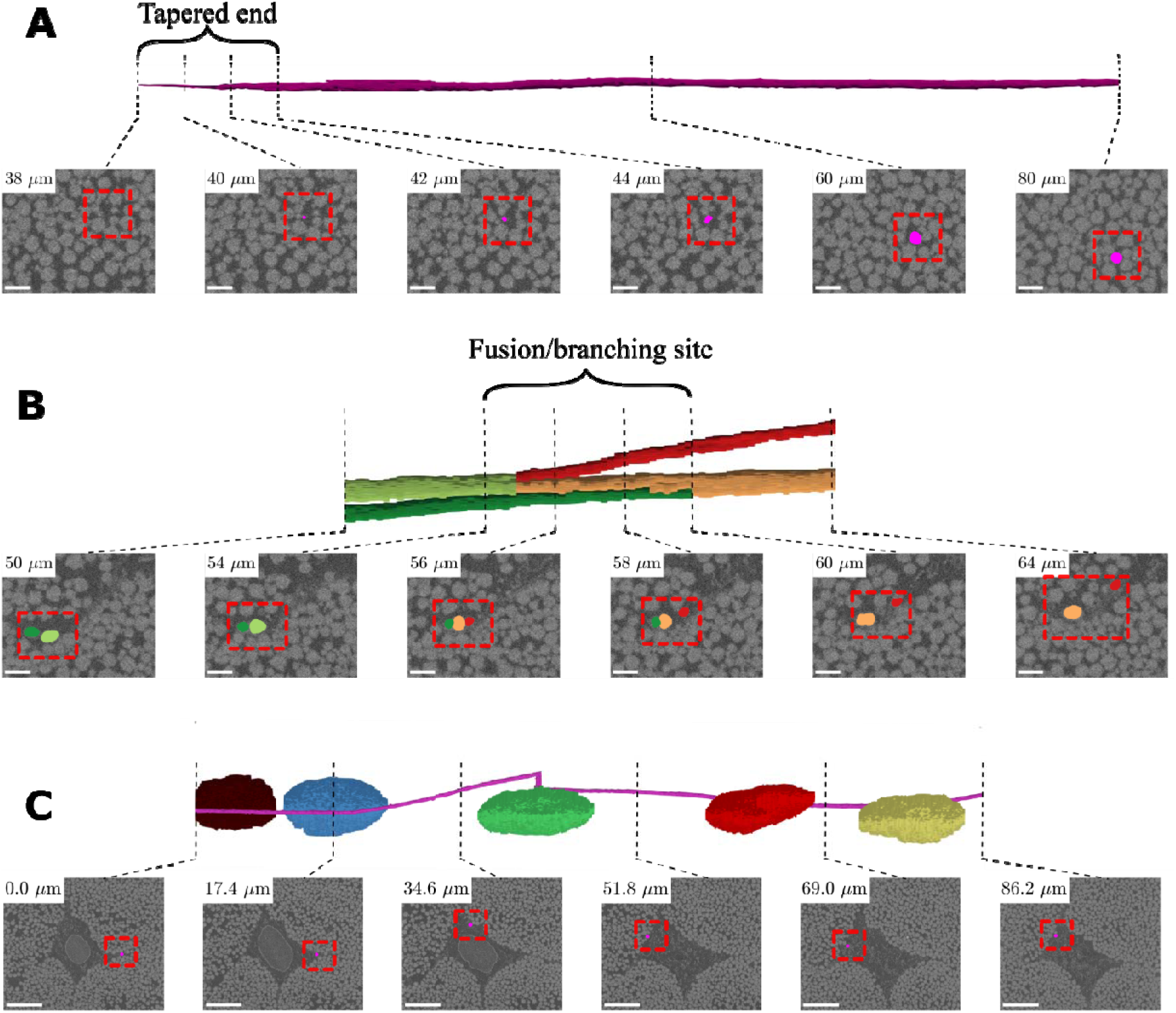
Three interesting but isolated structural features were observed: (A) tapered fibril end (B) fibril fusion/branching and (C) a helical fibril that wrapped around cells (only the nuclei are shown). For each feature, the first row includes the 3D segmentation, and the second row shows the segmentation mask overlaid on the SEM section image (dashed red box). In (A) and (B) scale bar = 1 *µm*, for (C) scale bar = 5 *µm*.

We made several isolated structural observations that have potential mechanical implications: tapered fibril end (Fig. 3A and Supplementary Video 1), fibril fusion/branching (Fig. 3B and Supplementary Video 2), and one fibril that wrapped around the cells (Fig. 3C and Supplementary Video 3). For the tapered fibril end, in the last ∼4 µm of the fibril length (scan depth of 38-42 µm) a reduction in fibril diameter was evident, where the fibril gradually fades away in the image sequence when approaching from the deeper scanned layers (Fig. 3A). At the fusion/branching site, a small fibril merges with a larger one, and in the subsequent scanned images the larger resulting fibril branches into two distinct ones. The fusion/branching site approximately spanned 6 µm, where at least two fibrils were not distinguishable (Fig. 3B). Another interesting feature was one fibril that made almost a full turn around the cells (Fig. 3C). These features were interesting but were isolated observations in the dataset.

When looking at the axial view of the fibrils, we observed several helical structures (Fig. 4 and Supplementary Video 4). In particular, many fibrils locally wrapped around each other, which contained two, three, or more fibrils with both left and right-handed helical configuration (Fig. 4 and Supplementary Video 5). Almost 50% (20 out of 42 fibril) of the fibrils that we segmented were in helical groups, although the sampling was not purely random. Of these helical fibrils, 13 fibrils had a right-handed twist and 7 were left-handed (Fig. 4B). These fibrils made an average of 2.2 ± 1.4 turns around each other along the scan-length (Fig. 4C), that also corresponds to an overall ∼39 ± 18 µm helical pitch (*λ*, the axial length of one full turn as described in Fig. 1F).

**Figure 4:**
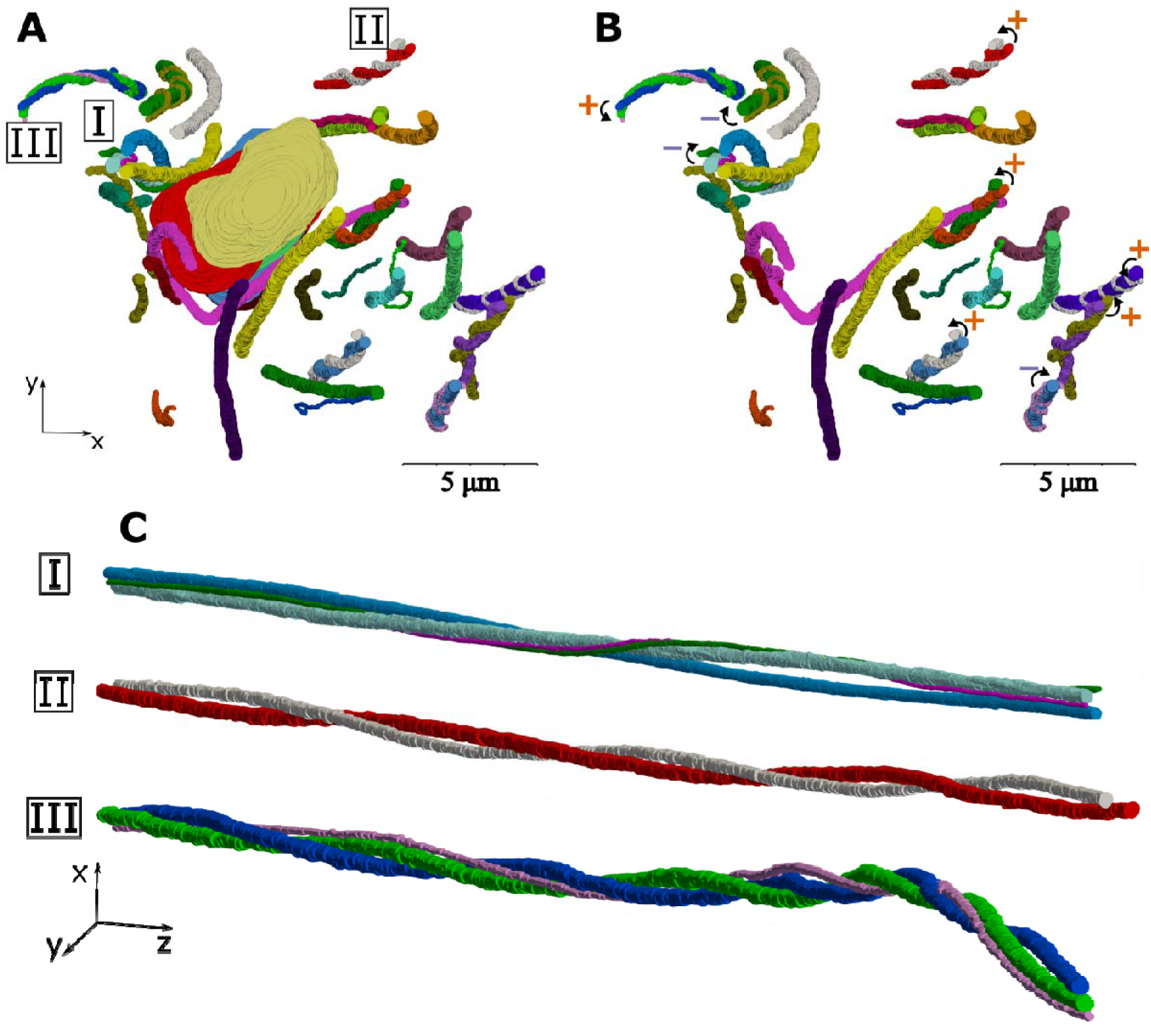
Many helical structures were observed in the axial view of the fibrils that are evident in the figures (A) with and (B) without the cell nuclei. The helical fibril groups have both left (marked with + ccw curve) and right-handed (marked with – cw curve) twist. (C) Shows three examples of the groups of twisting fibrils with pitch ranging from 20 um to 86 *µm*.

### Finite element simulations of the helical fibrils

We used a finite element model to test our hypothesis that frictional contact between helically wrapped fibrils can transfer stress (load) between fibrils without a need for a mediating matrix. The stress transfer was proportional to the fibril’s tensile modulus (Fig. 5A). Further parametric studies indicated that there is no change to the transferred stress with a change in Poisson’s ratio (*v*) (Fig. 5B). As expected, the transferred load increased with an increase of frictional coefficient (*µ*) (Fig. 5C). Our results show that the load transfer decreased with an increase in pitch (*λ*) (Fig. 5D).

**Figure 5:**
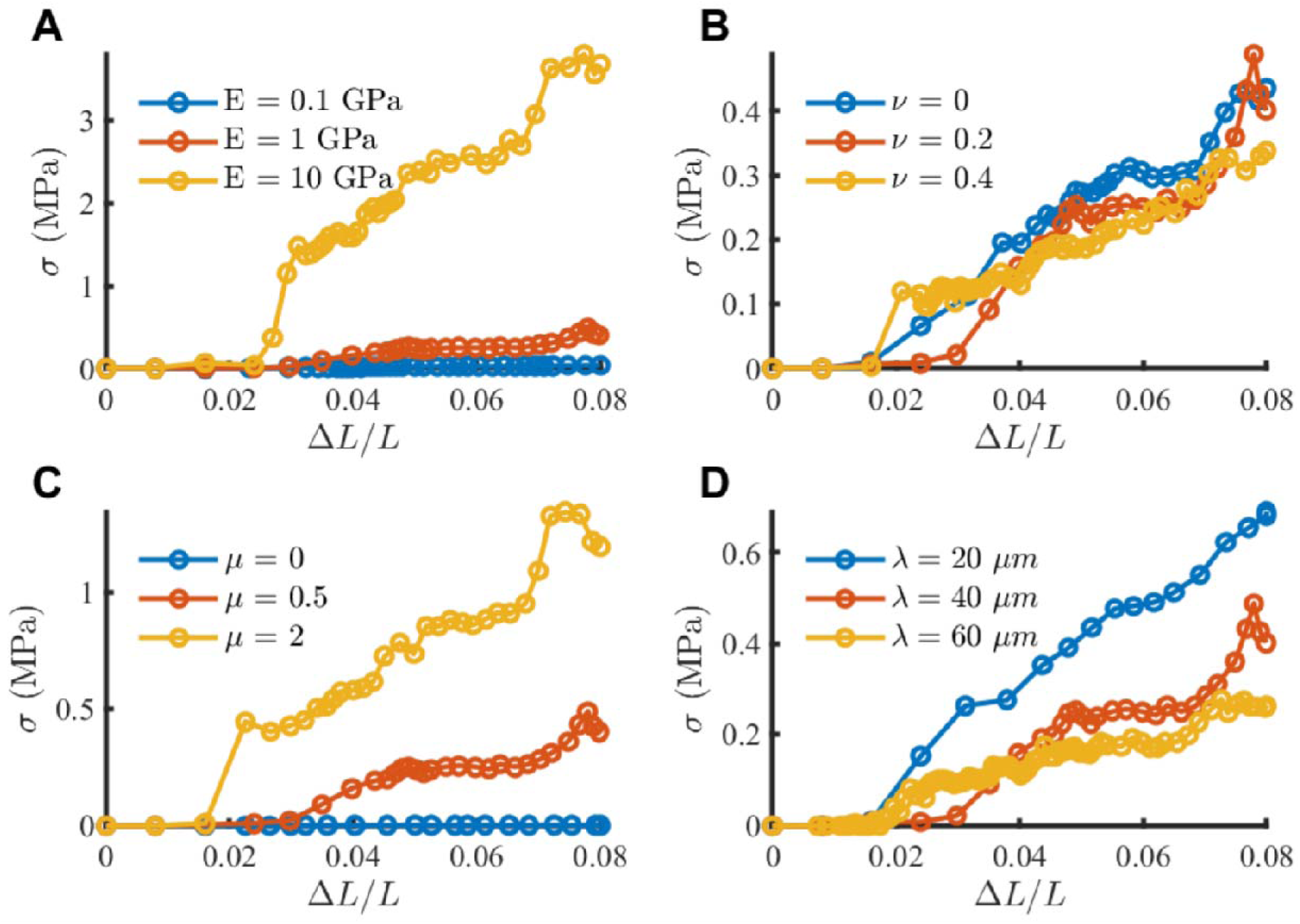
Finite element analysis of stress transfer due to friction between helically grouped fibrils. Base material properties *E* = 1 GPa, *v* = 0.2, *µ* = 0.5, *λ* = 40 *µm*. The transferred stress (A) increases with fibril modulus *E*, and (B) is insensitive to Poisson’s ratio *v*, (C) increases with frictional coefficient *µ*, and (D) decreases with increasing helix pitch *λ*.

To evaluate the spatial distribution of stress and deformation along the length of each fibril and its dependence on the friction coefficient, we plotted the axial stress and axial displacement (*ū*, Eq. 4) at the end of loading (Fig. 6). The fibril stress was zero with no friction (*µ* = 0, Fig. 6B) and it increased with higher friction coefficient (Fig. 6C and D). The axial stress varied linearly along the length of the fibril, increasing with distance from the free boundary and the stress in each fibril was the mirror image of the stress in the other one (Fig. 6A-D), which is in accordance with the static equilibrium condition. Similarly, for the fibril deformation, in the zero friction case, there was no axial deformation, hence the fibrils slid freely (Fig. 6E). When friction was increased, the induced deformation also increased, showing a plateau at the free ends indicating no strain, which confirms the stress-free boundary condition imposed on the model (Fig. 6F and G).

**Figure 6:**
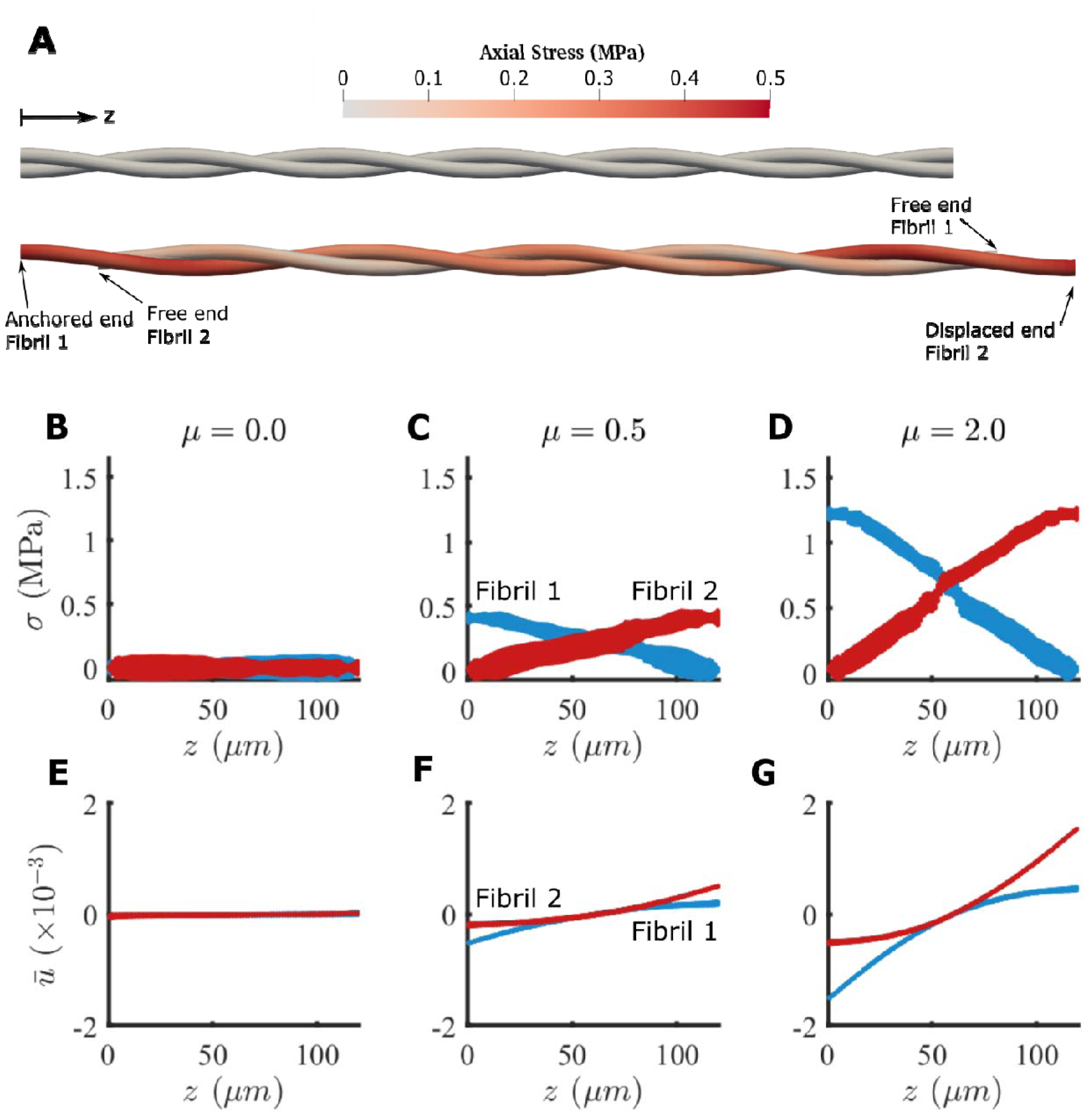
Finite element simulation of two helically wrapped fibrils. (A) Axial stress distribution in the fibrils for the undeformed (top) and deformed (bottom) states for *µ* = 0.5. The geometrical scale is compacted 10 times in the axial direction for clarity. (B–D) Stress along the length of the fibrils, and (E–G) normalized displacement (*ū*, Eq. 4) along the length of the fibrils for various choices of friction coefficient *µ*. With no friction (*µ* = 0) there was no *σ* or *ū* (B, E). As the frictional coefficient increases, the stress and the deformation increases for *µ* = 0.5 (C, F) and for *µ* = 2 (D,G). The model parameters are *E* = 1 GPa, *v* = 0.2, *λ* = 40 *µm*.

## Discussion

In this study, we visualized the microscale structure of tendon fibrils in three dimensions using serial block-face scanning electron microscopy (SBF-SEM) and studied the mechanical implications of helical fibrils as a mechanism for interfibrillar load transfer using FE analysis. We found that tendon fibrils are not purely parallel structures and there are many helical fibrils that wrap around each other and in groups. Our FE analysis indicated that, in addition to other potential mechanisms of load transfer (interfibrillar matrix and fusion/branching of smaller fibrils), the helical fibrils can also mediate load transfer through frictional mechanical contact.

### Microstructure of fibrils and SBF-SEM

We observed helical structures that twisted around each other (Fig. 4). The existence of helical fibrils was previously reported (Bozec et al., 2007; Orgel et al., 2006), and were suggested to be left-handed (Franchi et al., 2010); however, our findings showed a similar number of the left-handed and right-handed helical fibrils (Fig. 4B). The formation of the helical structure has been explained based on a fibripositor model of fibril assembly (Kadler et al., 1996; Kalson et al., 2015) and the helicity of subfibrillar structures (Bozec et al., 2007). The helical fibril structures explain the rotation of tendon in tension (Buchanan et al., 2017). Additionally, they have been used as an explanation for the macro-scale mechanical behavior of tendon, in particular for the low stiffness at small deformations and the large tensile Poisson’s ratio (Reese et al., 2010; Thorpe et al., 2013). The grouping of fibrils into helices can mediate load transfer by inducing frictional contact between fibrils during axial loading.

We observed some interesting but isolated features that can have implications for load transfer and loading on cells. For load transfer, we observed a tapered fibril end and fusion/branching, which agree with previous findings (Svensson et al., 2017), with the difference that the tapered end in this study was observed in a “straight” fibril in contrast to the hairpin shape in the previous study. As previously suggested by Szczesny and co-workers (Szczesny et al., 2017), an instance of fusion/branching, such as the one described above, could mediate load transfer between fibrils via direct physical connection. A free fibril end like the tapered end can also mediate load transfer by allowing microscale sliding and shear stress, which is a well-documented phenomenon in experimental studies (Lee et al., 2017; Szczesny and Elliott, 2014a; Thorpe et al., 2013). Additionally, we observed a fibril that wrapped around the cells, which indicates that during an axial loading, lateral compression can be exerted on the cells affecting tendon mechanotransduction (Lavagnino et al., 2015).

The fibril structure assessment was subject to some limitations. The SBF-SEM technique generates large 3D datasets and segmentation is consequently difficult compared to 2D SEM, creating practical limitations in data collection and analysis (Svensson et al., 2017). Although we consequently chose to do in-depth manual segmentation for only one sample, we performed several scans of additional fascicles, and these scans showed similar structural features. Additionally, although the helical fibrils were easily observed and represented approximately half of the total fibrils in the image sequence, future work using 3D automated segmentation will be needed to calculate the frequency of helical fibril. A separate limitation is that our observations only included a region near the tenocytes, and other regions further away from the cells might have a different structure. Furthermore, we only scanned the tail tendon, which is a low-stress tendon; further investigation is needed to confirm the existence of helical fibril groups in other tendons.

### Finite element simulations of the helical fibrils

Interfibrillar load transfer by friction in a helical contact does not require interfibrillar matrix bonding, but the sum of its contribution over many fibrils and its relative magnitude in comparison to other load transfer mechanisms such as chemical bonding remains unknown. Here, the estimated magnitude of the transferred fibril stress (∼0.2 - 4 MPa) was low compared to the ultimate stress of the fibrils—typically in the range of 90 MPa (Liu et al., 2016). The large variation in mechanical parameters of fibrils (Table 1) and the unknown accumulation among hundreds of fibrils hinders accurate calculation of the total load transferred in situ by the interfibrillar friction mechanism. Of particular importance, the fibril-on-fibril friction coefficient is unavailable. Thus, more experimental measurements of single fibrils and groups of fibrils are needed for more accurate estimations.

The variations in fibril stress transfer and displacement observed in the parametric FE sensitivity analysis make physical sense (Fig. 6). In particular, the dependence of transferred stress with modulus and frictional coefficient are expected based on Hooke’s law and Coulomb’s friction law, respectively, and they were shown using our FE model (Fig. 5A and C). Note, however, that while this studies used the relatively simplified model of friction contact, additional mechanisms, such as hydrodynamic friction, molecular asperity, and nonlinear dependence of friction coefficient to normal traction and other nano- and micro-scale tribological effects may be at interplay to induce load transfer between and within fibrils in the physiological system (Bhushan, 1999; Eyre et al., 2008; Sutcliffe et al., 1978; Ward et al., 2015). The decrease in reaction force by increasing the pitch of the helix, which is shown in Fig. 5D, can be explained by a reduction in lateral compression (normal force) on the fibrils as the pitch angle decreases and the fibrils become more parallel. The lack of dependence on Poisson’s ratio was an unexpected observation that might be due to the high aspect ratio of the fibril’s geometry or due to the isotropy of the constitutive relation (Fig. 5B). Large Poisson’s ratio in tension (*v*∼2) have been reported for individual fibrils (Wells et al., 2015), which is greater than the isotropic limit of *v* = 0.5. To include Poisson’s ratio’s effect on the interfibrillar load transfer, anisotropy of the isolated fibril material may be needed to be incorporated. The curves in Fig. 5 were not smooth, which is potentially related to the nature of frictional contact that can switch between slip and stick conditions and cause jitter (imagine moving heavy house furniture on ceramic), however the trends are clear, and this does not affect the outcome of the sensitivity analysis.

Interfibrillar friction causes a gradient of axial stress in the fibril. In the simulated cases (Fig 6), the spatial distribution of stress variation and deformation match the distributions of stress and deformation based on established shear-lag theories as applied to tendon fibrils (Szczesny and Elliott, 2014b). To further put this study in the context of prior research on theory of wire ropes, it should be mentioned that, unlike helical ropes such as elevator and bridge cables, which are continuous strands with only a small and local effect of friction (Costello, 1997; Jiang et al., 1999); in tissue, experimental observations demonstrate significant mircoscale sliding that mediates load transfer. Therefore, while there remains open questions regarding the physical existence and frequency of free fibril ends in mature tendon (Provenzano and Vanderby, 2006; Svensson et al., 2017), experimental tensile data suggests there are “effective” free fibril ends that allow microscale sliding (Szczesny et al., 2015). It is possible that fibril ends may exist in the form of tapered ends (such as Fig. 3A) or as physically weak links along the fibril length that can act as an “effective end” due to its high compliance (Veres et al., 2014). Further investigations are required to explore these microstructures and their contribution to interfibrillar sliding and shear load transfer. Regardless of how the effective fibril ends occur, the experimental evidence of microscale sliding and shear load transfer supports our FE model based on friction and its inherent boundary condition assumption for free fibril ends.

In conclusion, we used SBF-SEM to visualize the three-dimensional microscale tendon structure of the fibrillar network, and used FE analysis to demonstrate that helically arranged fibrils can have the mechanical function of frictional interfibrillar load transfer. Interfibrillar friction should be considered as another potential mechanism for interfibrillar load transfer, in addition to the previously postulated mechanisms of interfibrillar matrix shear and direct load transfer through fibril junctions. This study shows that a combined approach of SBF-SEM imaging and FE modeling is a powerful tool to study structure-mechanics relationships in tendon microstructure.

## Supporting information

Supplementary video 1: The animated mask for the tapered fibril end.

Supplementary video 2: The animated mask for the fibril fusion/branching.

Supplementary video 3: The animated mask for the fibril that wraps around the cells.

Supplementary video 4: The axial view of the in-plane trajectory of the fibril centroid

Supplementary video 5: 3D circular reconstruction of the fibril segmentation, showing the left and right-handed helical fibril groups.

## Acknowledgments

We would like to acknowledge Bioimaging center in the Delaware Biotechnology Institute (DBI) and the Thermo Fisher Scientific Company for SBF-SEM imaging. The funding for this study was made available from NIH-NIBIB grant R01-EB002425. Microscopy access was supported by grants from the NIH-NIGMS (P20 GM103446), the NSF (IIA-1301765) and the State of Delaware. The content is solely the responsibility of the authors and does not necessarily represent the official views of the NIH.

## Appendix: Supplementary dataset

The image stack for SBF-SEM images are available at (Safa et al., 2019).

